# Suboptimal eye movements for seeing fine details

**DOI:** 10.1101/220319

**Authors:** Mehmet N. Ağaoğlu, Christy K. Sheehy, Pavan Tiruveedhula, Austin Roorda, Susana T. L. Chung

## Abstract

Human eyes are never stable, even during attempts of maintaining gaze on a visual target. Considering transient response characteristics of retinal ganglion cells, a certain amount of motion of the eyes is required to efficiently encode information and to prevent neural adaptation. However, excessive motion of the eyes leads to insufficient exposure to the stimuli which creates blur and reduces visual acuity. Normal miniature eye movements fall in between these extremes but it is unclear if they are optimally tuned for seeing fine spatial details. We used a state-of-the-art retinal imaging technique with eye tracking to address this question. We sought to determine the optimal gain (stimulus/eye motion ratio) that corresponds to maximum performance in an orientation discrimination task performed at the fovea. We found that miniature eye movements are tuned, but may not be optimal, for seeing fine spatial details.

## Introduction

We make large, rapid, voluntary eye movements – saccades, to redirect our gaze to accomplish numerous visual tasks (e.g., searching for an object, reading a book, etc.) (Kowler, 2011). This is to form a fine-grained representation of the external world by taking advantage of a part of the retina – the fovea, which has the highest spatial resolution. However, our eyes are always in motion between epochs of saccades, even when we try to maintain our gaze on an object. Miniature eye movements that we make during fixation are often referred to as fixational eye movements (FEM). Different types of FEM have been identified depending on their spatiotemporal characteristics (Martinez-Conde, Macknik, & Hubel, 2004; Rucci & Poletti, 2015). Microsaccades are small jerky eye movements, and share similar peak velocity-amplitude dynamics as larger saccades (Otero-Millan, Troncoso, Macknik, Serrano-Pedraza, & Martinez-Conde, 2008). Drifts are relatively slower and smoother but rather erratic eye movements that occur between (micro) saccades, and have been usually modeled as various types of random walk or Brownian motion (Burak, Rokni, Meister, & Sompolinsky, 2010; Engbert, Mergenthaler, Sinn, & Pikovsky, 2011; Rucci, Iovin, Poletti, & Santini, 2007). Lastly, tremors are usually defined as very low-amplitude and high-frequency oscillatory movements that are superimposed on drifts (Ditchburn & Ginsborg, 1953; Ko, Snodderly, & Poletti, 2016; Ratliff & Riggs, 1950).

In addition to the non-uniform distribution of density and size of receptive fields of ganglion cells across the retina (Curcio, Sloan, Kalina, & Hendrickson, 1990; Dacey & Petersen, 1992; A. B. Watson, 2016), there is another unique property that differentiates our visual system from a computer vision system – neural adaptation. For instance, retinal ganglion cells (RGC) are most responsive to light transients and their responses decay with prolonged exposure (Benardete & Kaplan, 1997; Kaplan & Benardete, 1999). Although such a system is ideal for the detection of changes or movements that are crucial for survival in a natural setting, it comes with a consequence. It has been known that in the absence of FEM, visual perception fades away, and more so for small visual stimuli (Ditchburn & Ginsborg, 1952; Riggs, Ratliff, Cornsweet, & Cornsweet, 1953; Yarbus, 1967). This suggests that retinal image motion is essential for continuous and high-acuity vision. However, if FEM are too large or too fast, the light intensity defining a visual stimulus will spread over a large population of cells each with an insufficient exposure to the stimulus, resulting in nothing but smeared, ghost-like impressions. Therefore, there must be an optimum movement between these two extremes whereby visual perception is maximized such that the ability to perform a visual task is highest at that particular movement.

Previous research showed that naturally occurring or artificially induced irregular and continuous retinal image drifts help in seeing fine spatial details (Ratnam, Harmening, & Roorda, 2017; Rucci et al., 2007). Likewise, naturally occurring microsaccades or sudden jumps of stimuli are known to counteract visual fading (Costela, McCamy, Macknik, Otero-Millan, & Martinez-Conde, 2013; Martinez-Conde, Otero-Millan, & Macknik, 2013!), and help redirect our gaze to compensate for the non-homogeneous acuity within the fovea (Poletti, Listorti, & Rucci, 2013). If there is a causal relationship between FEM and visual perception, the latter should show a “tuning” function with different levels of the former, analogous to orientation tuning of cells in the early visual areas where firing rate of a given cell is continuously modulated by how close the orientation of a stimulus in its receptive field to its “preferred orientation.” In other words, direct manipulation of FEM, or the way retinal image moves as a function of FEM should result in systematic changes in visual perception, where maximum performance in a visual task would be obtained at the preferred or optimal FEM. From an evolutionary point of view, FEM in normal vision can be thought as optimally tuned for seeing fine details at the fovea. Here, we explicitly tested this hypothesis. Our results show that normal FEM are tuned, but not quite optimal, for fine discrimination at the fovea. We also found that within the range of spatial frequencies where the human visual system has highest contrast sensitivity, this relationship disappears suggesting a higher tolerance for retinal image motion for coarse visual structures.

## Methods

### Participants

Seven human subjects (including the first author, S1) with normal or corrected-to-normal vision (20/20 or better in each eye) participated in the study. All seven subjects took part in Experiment 1. Three of the seven subjects participated in Experiment 2. All subjects, except the first author, were naïve as to the purpose and the details of the experiments. All subjects gave written informed consent prior to the experiments. All experimental procedures followed the principles put forth by the Declaration of Helsinki, and were approved by the Institutional Review Board at the University of California, Berkeley.

### Apparatus

For stimulus delivery and eye tracking, we used a custom-built tracking scanning laser ophthalmoscope (TSLO) (Sheehy et al., 2012). The TSLO has a diffraction-limited optical design, provides high-fidelity imaging of the retina, and more importantly, offers online tracking of eye movements. For the experiments presented here, we used a 10 x 10 deg^2^ (512 x 512 pixels^2^) field of view (FOV), which yielded a pixel size of 1.17 arcmin. A large FOV enabled us to capture videos with rich retinal structure which, in turn, allowed accurate image-based eye tracking and stimulus delivery. The horizontal scanner operates at 16 kHz whereas the vertical scanner operates at 1/512 of this rate to record full frames at 30 frames per sec. An 840 nm super luminescent diode with a 50 nm bandwidth was used to scan the retina. Visual stimuli were delivered by manipulating the laser beam via an acousto-optic modulator with the output controlled by a 14-bit digital to analog converter. Therefore, the stimuli had a negative contrast on the dim red raster created by the scanner (i.e., appeared black on a red background). Details of online eye movement tracking have been reported elsewhere (Arathorn et al., 2007; Mulligan, 1997; Yang, Arathorn, Tiruveedhula, Vogel, & Roorda, 2010). Briefly, each frame was broken into 32 horizontal strips (16 x 512 pixel^2^) and each strip was cross-correlated with a reference frame acquired earlier. The horizontal and vertical shifts required to match a strip to the reference frame represent a measure of the relative motion of the eye. This method results in an eye movement sampling rate of 960 Hz. These computations occur in near real-time (with 2.5±0.5 ms delay) and allows accurate stimulus delivery at specific retinal locations.

### Stimuli and Procedures

The task was to report the orientation of a sinusoidal grating from vertical (2AFC, clockwise or counterclockwise). The amount of tilt was ±45° from vertical and the spatial frequency of the grating was 12 cpd in Experiment 1 (**Fig. 1a**) and 3 cpd in Experiment 2. Prior to each experiment, the contrast of the grating was adjusted for each subject to yield ∼80% correct discrimination performance under the natural viewing condition (i.e., gain = 0). The size of the grating was 3 x 0.75 deg^2^. Horizontal edges were smoothed using a cosine profile. Mean luminance (as measured indirectly from laser power) of the grating was kept at ∼70% of the luminance of the scanning raster for all subjects, regardless of the contrast of the grating (**Fig. 1b**). All experiments were performed in a dark room, and subjects were dark adapted for about half an hour prior to any data collection. For some subjects (three out of seven), pupil of the imaged eye was dilated to maintain good retinal image quality throughout the session.

**Figure 1.**
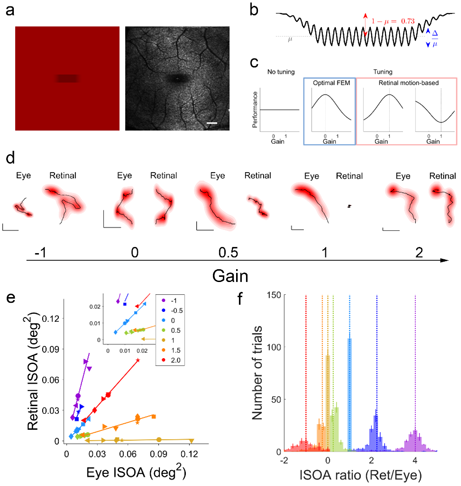
Manipulating the relationship between retinal image motion and eye motion with the TSLO. **(a)** An orientation discrimination task at the fovea. Subjects' view of a grating on the raster (left), and corresponding retinal image (right). Note that the stimulus is imprinted on the retinal image. **(b)** The luminance profile used to create grating patterns on the raster. The mean luminance of the grating was set to ∼70% of the background and the contrast of grating was adjusted for each subject. **(c)** Predictions from the no tuning (null) and tuning hypotheses. The panels with blue and red outlines show various ways tuning can occur. **(d)** Sample eye motion and retinal image motion traces (black lines) and corresponding probability densities (red clouds) for different gains. The horizontal and vertical lines in the lower left corner of each panel represent 0.1°. Dimensions were adjusted for clarity. **(e)** Retinal ISOA as a function of eye ISOA across gains in Experiment 1. Different colors represent different gains, and subjects are coded by different symbols. Inset shows a close-up view of data for smallest retinal/eye motion. **(f)** The distribution of retinal/eye ISOA ratios for different gains, averaged across seven subjects. Vertical dotted lines show theoretical ISOA ratios, i.e., assuming that eye tracking, stimulus delivery, and offline eye movement extraction were perfect. Error bars represent ±SEM (n=7). Color conventions for gains are identical across all figures.

Each trial started with a fixation cross (0.2 deg) presented at the center of the raster. The experimenter manually acquired a reference frame for online tracking at the start of each trial. Following a random delay (up to 1 sec), the stimulus was presented for 900 ms (flickered at 30 Hz). Subjects responded via a gamepad, had unlimited time to respond, and could have a break at any point during a block of trials. The set of gains used were −1, −0.5, 0, 0.5, 1, 1.5, and 2. Different gains were interleaved within a block of trials. All observers completed at least 100 trials per gain. Right after the completion of the main experiment and before data analyses, subject S4 was tested for a second run in Experiment 1 since her retinal image quality and head stability during the experiment was poor, causing retinal stabilization to fail in many of her trials. In the second run, we used finer steps of gain (from −0.25 to 1.25 in steps of 0.125) and subject S4 again ran at least 100 trials per gain. However, all results presented in the main text include only the data from the first run for S4. The results from both runs for S4 are shown in **Fig. 3**.

**Figure 3.**
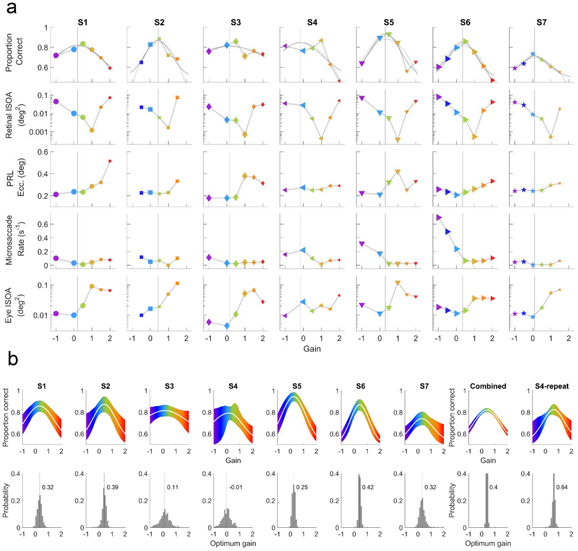
Individual results from Experiment 1. **(a)** Proportion correct and all mediators quantified in the present study, binned based on gain. **(b)** Bootstrapping tuning curve fits (top) and optimal gains (bottom) for each subject by using binary data (correct vs incorrect). White lines represent the fits corresponding to median parameters. Shaded regions (top) represent 2.5-97.5% percentiles of the bootstrapped distributions of fitted curves. For each panel, bootstrapping was done by resampling the individual trial data with replacement 1000 times. Vertical dashed lines (bottom) represents the median optimal gains. The data from second run of S4 were combined with the first run and analyzed together, and are shown here in the rightmost panels. The vertical axes in bottom panels in **(b)** are cropped to 0.4 for better visibility.

### Retinal video analysis

Retinal videos were analyzed offline for five main reasons. First, we sought to determine how well and where the stimulus was delivered on a trial-by-trial basis. Second, online eye tracking was performed by using raw retinal images which were corrupted by high frequency noise and low frequency luminance gradients. In order to get more accurate eye motion estimates relatively less dependent on changes in overall brightness and uniformity of retinal images across trials, one needs to perform several preprocessing steps on retinal images. To this end, we performed the following image processing steps before computing eye motion. Trimming, detection and removal of frames during which subjects blinked, extracting stimulus position and removal of the stimulus (replaced by random noise patterns whose statistics –mean and standard deviation, matched to the rest of the frame), gamma correction, bandpass filtering (for removal of high frequency noise and low frequency brightness gradients), and making a reference frame. Third, during online eye tracking, if the peak of normalized cross-correlation between a strip and the reference frame was below 0.3, possibly due to (i) bad image quality, (ii) excessive distortion of image features due to a rapid eye movement, (iii) insufficient amount of overlap between the strip and the reference frame due to large eye motion, or (iv) blinks, the stimulus was not delivered. By offline processing of retinal videos, we also sought to inspect each and every frame of retinal videos and discard the trials if the stimuli was delivered inaccurately or was not delivered at all in more than two frames per trial (note that with this criterion, trials where subjects blinked were also discarded). This procedure resulted in removal of 28.8% (1305/4525) and 8.7% (206/2353) of all trials in Experiment 1 and 2, respectively. Fourth, since the reference frames used for online tracking were basically snapshots of the retina taken manually by the experimenter, and since the eyes are almost never stationary, the reference frames themselves might have some distortions due to these motions.

By offline processing, we created a relatively motion-free reference frame for each and every trial separately in an iterative process. This process started by selecting one of the frames in a retinal video as the reference frame, and computing eye motion. After each iteration, a new reference frame was built by using computed eye motion and individual strips. We performed three iterations for each video, and the reference frames made in the last iteration were used for the final computation of eye motion. The strip height and sampling rate used for the final strip analysis were 25 pixels and 540 Hz, respectively. Fifth, during offline analysis, we could interpolate the cross-correlation maps around where the peak occurs to achieve subpixel resolution (one tenth of a pixel, 0.12 arcmin) in computing eye motion.

### Post-processing

Following strip analysis of individual videos, the computed eye motion traces were subjected to several post-processing steps. First, eye motion traces were “re-referenced” to a larger reference frame created by retinal videos recorded in a separate session where subjects were asked to fixate at different position on the scanning raster. This essentially allowed us to capture images from different part of the retina and tile them on a larger (“global”) reference frame. Re-referencing was needed since each and every video had a slightly different (“local”) reference frame (since reference frames were created for each video separately), and hence, the absolute values of the eye motion would differ across trials. This step is required also for computing the absolute retinal position of the stimulus across all trials for a given subject. Re-referencing was performed by adding a constant shift to the previously computed eye motion traces, where the amount of shift was defined as the position of the local reference frame on the global one. After re-referencing, eye motions were then converted to visual degrees, low-pass filtered (passband and stopband frequencies of 80 and 120 Hz, respectively, with 65 dB attenuation in the stopband), and median filtered (with a window of 9 samples, ∼17 ms) to reduce frame-rate artifacts (30 Hz noise and its harmonics). Filtered traces were used to compute retinal motion of the stimulus (defined as the difference between stimulus motion and eye motion). We quantified the amount of eye and retinal motion on a trial-by-trial basis as the 68% isoline area (Castet & Crossland, 2012) (referred to as ISOA in the main text), which corresponds to the area of the 0.68 cumulative probability isoline (**Fig. 1d**). This also roughly corresponds to the area covered by 68% percent of the motion samples. In the case of retinal motion, this metric quantifies the retinal area traversed by the stimulus in a given trial, and for eye motion, it represents the area of the raster (in world centered coordinates) over which the eye moved. The distribution of the ratio of retinal ISOA and eye ISOA reveals how well the stimulus was delivered in different conditions (see Fig. X). The theoretical ratio for a given gain is defined as

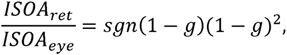

where *g* represents gain, and *sgn*(.) represents the signum function which was introduced to differentiate between gains that result in the same retinal motion magnitude but in opposite directions (**Fig. 1f**, and **Fig. 3c**).

We also computed the preferred retinal locus (PRL) of the stimulus for each trial. PRL was defined as the retinal location corresponding to the highest probability density of stimulus presence. The probability densities were computed by the “kernel density estimation via diffusion” method (Botev, Grotowski, & Kroese, 2010) with a slight modification. More specifically, the kernel bandwidth was set to one-sixth of the standard deviation of the eye (or retinal) motion (as in Kwon, Nandy, & Tjan, 2013). The median PRL in the trials where the gain was 0 (i.e., natural viewing) was taken as the location of the fovea and trial-to-trial PRL eccentricity was calculated with respect to this quantity. Finally, we identified microsaccades by using a median-based velocity threshold (Engbert & Kliegl, 2003). Eye motion traces from all trials were visually inspected to ensure that microsaccade detection was performed correctly.

The PRL estimated when gain is 0 reflects the true PRL. Since PRL is mostly determined by the position of the stimulus at the beginning of a trial (e.g., when gain is 1, the stimulus will stay at the start position), the estimated PRLs in other gain conditions do not necessarily reflect the preferences of the subjects. However, they demonstrate the idiosyncratic eye movements which govern the starting position of the stimulus. Nevertheless, to keep a consistent nomenclature, we used the term PRL across all conditions.

### Statistics

In order to test the tuning and no tuning hypotheses, we fit performance with a flat line and a quadratic polynomial (as well as a Gaussian, although polynomial and Gaussian fits produced almost identical results), and compared the adjusted R^2^ values as a metric of goodness of fit.

Due to foveal presentation of the stimuli, different gains led to different idiosyncratic oculomotor behaviors which could not be controlled during the experiments. We quantified several covarying factors such as retinal ISOA, eye ISOA, PRL eccentricity, and microsaccade rate. The exact choice of covarying factors was driven by the need to account for main retinal and eye movement-related metrics. It is possible to estimate retinal image velocity, acceleration, or components of retinal image motion parallel or perpendicular to the orientation of the gratings. It is also possible to quantify these metrics in multiple ways, such as by their mean, standard deviation, minimum, maximum, or any combination of these together. However, since most of these metrics are strongly related to each other, adding different variants of them does not add much explanatory power. In addition, eye position traces extracted from retinal videos tend to have frame-rate artifacts, i.e., more power than normal at temporal frequencies around the frame rate of the videos. The frame-rate artifact gets amplified for velocity and acceleration due to differentiation, and more importantly, the severity of the effect interacts with different gain conditions (due to changes in oculomotor behavior). Position estimates are not influenced as much and indirectly captures the effect of its derivatives.

In order to determine how much of the effect of gain on performance is mediated by the aforementioned covarying factors, we performed a linear-mixed effects regression-based mediation analysis (MacKinnon, Fairchild, & Fritz, 2007). We followed the commonly-used four-step approach suggested by Baron and Kenny (Baron & Kenny, 1986). The aim of this analysis was to determine whether or not gain had a significant direct effect on discrimination performance, even after taking into account the effects of mediators (i.e., significant covarying factors). Mediation analysis can be done in many ways depending on the causal relationship between the independent variable and mediators. When there are multiple mediators, say n, there are 2^(n!)^ possible ways of decomposing total effect size. In our case, n=4, this yields 16,777,216 possibilities (Daniel, De Stavola, Cousens, & Vansteelandt, 2015). Since finding the best way of organizing mediators to account for data is beyond the scope of the present work, we chose a simple case where all mediators were treated as independent factors that are directly modulated by only the independent variable, the gain, and they did not have interactions among each other nor with the gain. However, they were allowed to covary with subjects, and each had a fixed slope and a random intercept to account for individual differences in mediator values.

## Results

Using a tracking scanning laser ophthalmoscope (TSLO) (Sheehy et al., 2012), we presented seven human subjects with a high spatial frequency grating (12 cpd) for 900 ms while imaging their retina and tracking their eye movements in real time (**Fig. 1a, b**). We systematically manipulated the way retinal image motion and the actual FEM are related. The motion of the stimulus on the scanning raster was a function of the estimated eye motion times a *gain* factor (**Fig. 1d**). A gain of 0 means that the stimulus position remained fixed relative to the raster but slipped across the retina based on natural FEM. A gain of 1 means that the stimulus was stabilized on the retina, i.e., the retinal image motion due to FEM was completely cancelled out. A gain of 0.5 refers to partial stabilization, i.e., the stimulus moved only half as much as the eye motion. Assuming similar oculomotor behavior under different gains, a gain of −1 doubles the retinal slip of the stimulus compared to that under natural viewing (i.e., gain = 0), and a gain of 2 results in the same retinal slip but in the opposite direction of what would occur under natural viewing. Offline analyses of retinal videos for eye movement extraction revealed that eye tracking and stimulus delivery were performed with near-perfect accuracy (∼99%) for complete retinal stabilization (gain = 1). For gain conditions other than 0 and 1, there was some trial-to-trial variability in accuracy of stimulus delivery (**Fig. 1e, f**). Nevertheless, each gain condition resulted in a statistically distinct distribution of effective gains centered at the desired gain (**Fig. 1e, f**). We measured subjects' ability to discriminate the direction of the grating's orientation from vertical under different gain conditions. If FEM are not tuned for fine discrimination at the fovea, then performance should not depend on gain (the null hypothesis, **Fig. 1c**). On the other hand, if the retinal image motion due to FEM is tightly tuned for fine discrimination, performance should manifest a non-monotonic relationship with gain, where a particular value of gain results in the best (or worst) performance (**Fig. 1c**). The tuning hypothesis can hold true in various ways. If FEM are optimal for fine discrimination at the fovea, then visual performance should peak at the gain of 0. Alternatively, retinal image motion might be the primary determinant of visual performance. If retinal image motion is always detrimental for seeing, visual performance should be highest at the gain of 1. Retinal motion might also be beneficial regardless of the underlying FEM. If that is the case, the lowest discrimination performance should occur when the stimulus is fully stabilized on the retina.

We found that orientation discrimination performance is tuned to gain (**Fig. 2a, b**). Averaged across subjects, the peak performance occurred at a gain of 0.43 (95% confidence intervals: 0.12, 0.74), suggesting that partially reducing the effects of FEM is actually helpful in seeing fine spatial details. Results were similar when data from each subject were fitted separately (polynomial: t_6_=3.165, p=0.019; Gaussian: t_6_=2.600, p=0.041) (**Fig. 2b** and **Fig. 3**). The exact choice of the tuning model (a quadratic polynomial or a Gaussian) did not matter (paired t-test: t_12_=3.165, p=0.019). Bootstrapping tuning curve fits to binary data (correct vs incorrect) also revealed no optimality in six of the seven subjects (**Fig. 3**). These results suggest that FEM are tuned but not optimal for fine discrimination at the fovea, at least within the range of parameters investigated here.

**Figure 2.**
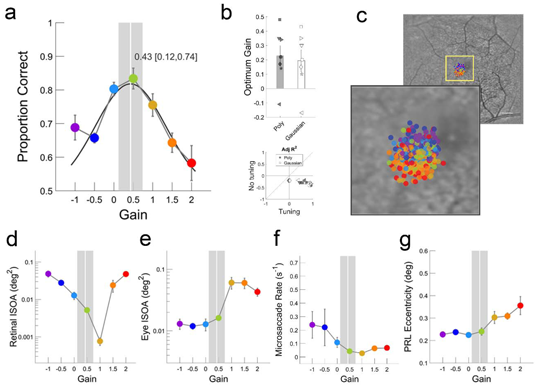
FEM are tuned, but not optimal, for fine discrimination at the fovea. **(a)** Proportion correct as a function of gain, averaged across subjects, in Experiment 1. A Gaussian tuning function was fit to all data (black curve) to estimate the optimum gain. Vertical white line represents optimal gain defined as the gain corresponding to the peak of the Gaussian. Shaded regions represent 95% confidence intervals of the optimum gain. **(b)** (Top) Average optimal gains based on individual tuning function fits along with individual optimal gains. A quadratic polynomial and a Gaussian tuning function resulted in statistically indistinguishable optimal gains. (Bottom) To compare the “no tuning” and “tuning” hypotheses in terms of how well they can explain our data, we computed Adjusted R^2^ metric for the constant model and tuning models (a quadratic polynomial or a Gaussian), respectively. For all subjects, tuning models performed better. **(c)** (Top-right) the distribution of preferred retinal locus (PRL) across trials for one representative subject. Each symbol represents one trial. (Bottom-left) A close-up view of the central ∼2.5° part of the retina. Note the systematic change in PRLs across gains. **(d, e)** Retinal image motion and eye motion ISOA as a function of gain. **(f, g)** Microsaccade rates and PRL eccentricity across gains. Optimal gain and confidence intervals in (a) are replotted in (d, e, f, and g). Error bars represent ±SEM (n=7).

In order to check whether or not the tuning between performance and gain is limited only to fine discrimination tasks, we repeated the experiment with a spatial frequency (3 cpd) at which the human visual system has highest contrast sensitivity for static displays(Kelly, 1977). The hypothesis was that the retinal jitter due to FEM causes much less modulations in retinal ganglion cells with low spatial frequency stimuli, therefore, gain manipulations should result in minimal or no change in performance. We found no effect of gain on performance (**Fig. 4a**), despite the retinal image motion varied over two log units across conditions (**Fig. 4d**). Different performance trends with gain in Experiments 1 and 2 cannot be explained by differences in retinal motion, eye motion, microsaccade rate, or PRL eccentricity (**Fig. 2d-g** vs **Fig. 4d-g**).

**Figure 4.**
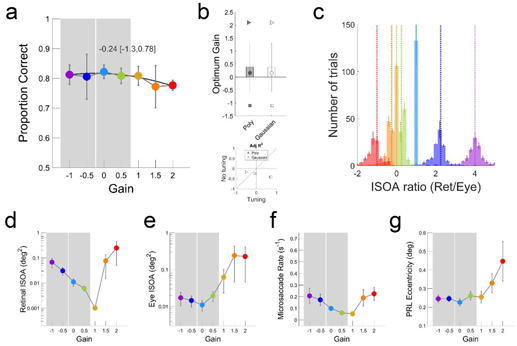
FEM are not tuned for coarse discrimination at the fovea. **(a)** Proportion correct as a function of gain in Experiment 2. **(b)** Average optimal gains based on individual tuning function fits along with individual optimal gains. **(c)** The distribution of retinal/eye motion ISOA in Experiment 2. **(d, e)** Retinal image motion and eye motion ISOA as a function of gain. **(f, g)** Microsaccade rates and PRL eccentricity across gains. Optimal gain and confidence intervals in **(a)** are replotted in (d, e, f and g). Error bars represent ±SEM (n=7). Conventions are as in **Figure 2.**

Next, we sought to determine what drives the strong dependency between performance and gain with high spatial frequency gratings. If gain manipulation only modulates the retinal image motion, then the answer would simply be retinal image motion, assuming no interference from extra-retinal mechanisms. The approach taken in most retinal stabilization studies in the literature implicitly assumes that gain manipulation only results in changes in retinal image motion. In other words, retinal image motion is considered as the one and only mediator of performance. However, we found that gain modulates multiple mediators. We computed two-dimensional probability density of stimulus locations on the retina and eye positions on the raster (e.g., **Fig. 2d**), and quantified, on a trial-by-trial basis, the extent of retinal image motion and eye motion by the isoline area (ISOA) containing roughly 68% of the retinal/eye motion traces (**Fig. 2d, e** and **Fig. 3**). As expected, the minimum retinal ISOA occurred when the stimulus was stabilized on the retina (i.e., gain = 1) but the pattern of changes in retinal ISOA as a function of gain revealed an asymmetric “V” shape around the gain of 1 (**Fig. 2d**). This asymmetry can be explained by differences in oculomotor behavior of subjects across different gains. More specifically, consistent with previous literature (Poletti, Listorti, & Rucci, 2010), subjects made smooth pursuit-like eye movements for gains of 1 and larger, which resulted in larger eye ISOAs (**Fig. 2e**). This change in behavior occurred as soon as the retinal slip is no longer in a direction that is consistent with eye motion, in line with recent perceptual observations (Arathorn, Stevenson, Yang, Tiruveedhula, & Roorda, 2013). In addition, subjects made slightly more microsaccades for negative gains where retinal image motion is amplified (**Fig. 2f** and **Fig. 3**). In addition, although each trial started with a fixation cross at the center of the raster, the preferred retinal locus (PRL) during grating presentation, defined here as the retinal location corresponding to peak probability density of retinal stimulus locations, also changed with gain (**Fig. 2c, f, g** and **Fig. 3**). To determine what really drives the relationship between gain and performance, one must take these mediators into account. In a regression-based mediation analysis following the most commonly used four-step approach (Baron & Kenny, 1986), we found that (i) gain has a significant effect on performance, (ii) gain significantly modulated all four mediators (retinal ISOA, eye ISOA, microsaccade rate, and PRL eccentricity), (iii) all mediators individually, with the exception of microsaccade rate, are significant predictors of performance, (iv) gain remains a significant predictor of performance even when the effects of all significant mediators are taken into account (**Fig. 5**). In order to determine whether or not mediators can account for the data as well as gain by itself, we performed a series of linear-mixed effects regression analyses (**Fig. 6**). In terms of explained variance and log-likelihood, where number of factors is not penalized, several purely mediator-based models could surpass the models based on gain only, suggesting that mediators identified here might fully account for how gain modulates performance. However, as the Bayes Information Criterion (BIC) differences show, none of the mediator-based models could outperform the simple model that is based only on gain. Through additional regression analyses and model comparisons using BIC, we confirmed that performance cannot be fully accounted by mediators alone (**Fig. 6**).

**Figure 5.**
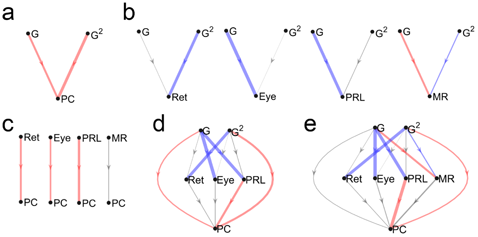
Teasing apart contributions of different mediators. **(a)** The first step in mediation analysis is to establish a significant relationship between gain (G) and proportion correct (PC). Since the tuning hypothesis predicts a quadratic relationship between G and PC, we included the G^2^ in our regression analyses. **(b)** Second, whether or not gain is a significant predictor of each covarying factor (Ret: retinal ISOA, Eye: eye ISOA, PRL: PRL eccentricity, MR: microsaccade rate) is established. **(c)** The third step tests separately for a significant effect each mediator on performance. **(d)** Finally, gain and mediators with a significant correlation on performance are used to explain performance. Red and blue colors represent statistically significant negative and positive effects whereas gray lines represent insignificant relationships. The final model in **(d)** shows that even when all significant mediators are taken into account, gain still has a significant effect on performance. **(e)** When all mediators are included, regardless of the outcome of **(c)**, gain remains to be a significant factor. Thickness of each line represents the absolute value of the standardized effect size.

**Figure 6.**
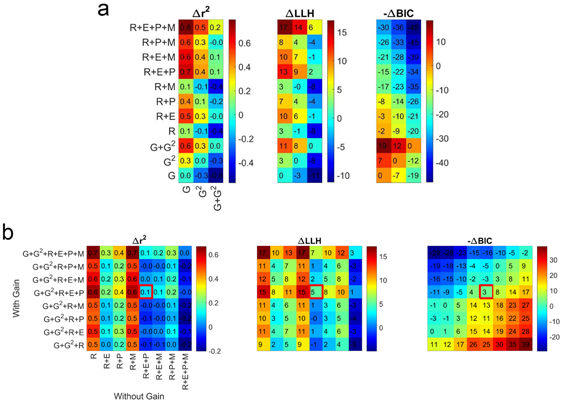
Contributions of gain and mediators in explaining variance. **(a)** Change in (left) explained variance, (middle) log-likelihood, and (right) BIC with addition of mediators. Note that the sign of ΔBIC is flipped so that red color represents superiority of a model on the vertical axis with respect to another one on the horizontal axis. G: Gain, R: retinal ISOA, E: eye ISOA, P: PRL eccentricity, M: microsaccade rate. **(b)** Can mediators fully account for the effects of gain? Here, we explicitly tested whether having gain in addition to mediators improve statistical models substantially. The right diagonal in each panel represents the exact contribution of the gain term. The red squares represent the final model in the mediation analysis (G+G^2^+R+E+P) (Fig. 5d). In general, adding gain was helpful only when there are three or less mediators in the regression model.

## Discussion

"Tuning" refers to a relationship between an independent variable and an outcome measure, where a certain level of the former is more preferable than others. Optimality in this context refers to achieving the best possible outcome in the face of several antagonist factors. Throughout the vast literature on FEM and visual perception, the word “optimal” has been used quite liberally in regard to spatiotemporal properties of FEM (Ahissar & Arieli, 2012; Cornsweet, 1956; Ditchburn, Fender, & Mayne, 1959; Gerrits & Vendrik, 1970; Kuang, Poletti, Victor, & Rucci, 2012; Martinez-Conde et al., 2004; Skavenski, Hansen, Steinman, & Winterson, 1979) although there has never been an explicit test for addressing it. Here, we tested whether visual performance in a fine orientation discrimination task would show tuning as a function of the relationship between the retinal image motion and actual eye movements. We found strong tuning for a fine-detail discrimination task (Experiment 1) but not for a coarse discrimination task (Experiment 2). The absence of tuning in Experiment 2, despite up to a two log-unit change in retinal motion across conditions, suggests a very high tolerance for motion. Surprisingly, the optimal gain in Experiment 1 was obtained at a gain value between 0 and 1, suggesting that partially compensating for FEM can be beneficial.

Our results might seem inconsistent with previous reports where complete retinal stabilization resulted in impaired discrimination performance (Ratnam et al., 2017; Rucci et al., 2007). A simple interpolation between the two extremes suggests a monotonic impairment in visual performance with better compensation for FEM. This apparent inconsistency may not be real. First, it is technically possible to get impaired performance with complete stabilization *and* a nonzero optimal gain at the same time (which was the case for five out of seven subjects, **Fig. 3**). Second, none of the existing studies explored the range of gains used here for discrimination tasks at the fovea. In addition, in previous studies, fading that resulted from retinal stabilization was quantified by threshold elevations, but the degree to which fading occurs depends on many variables such as stimulus duration, size, contrast, eccentricity, equipment used, etc. (Coppola & Purves, 1996; Kelly, 1979a, 1979b; Riggs et al., 1953; Riggs & Tulunay, 1959). Early studies on retinal stabilization used small stimuli extending only a few arcmin, and was closely surrounded by other visual cues coming from the apparatus (Ditchburn & Ginsborg, 1953; Riggs et al., 1953; Yarbus, 1967). More recent studies used foveally presented gratings extending several degrees of visual angle far from display boundaries (Poletti et al., 2013; Rucci et al., 2007), or parafoveally presented diffraction-limited stimuli covering only a few cones within a visible raster covering 1-1.3 deg (Ratnam et al., 2017). The paradigm used here was somewhere in between; we presented through natural optics of the eye a grating that covers the fovea and is situated within a visible raster covering 10 deg. Therefore, it is possible to make qualitative comparisons across aforementioned studies, however, it is not feasible to extrapolate previous studies to the conditions investigated here.

Our results are highly consistent with recent theoretical work that has been successfully used to account for performance impairment due to retinal stabilization (Anderson, Olshausen, Ratnam, & Roorda, 2017; Burak et al., 2010; Pitkow, Sompolinsky, & Meister, 2007). According to this framework, there are two distinct mechanisms that work in tandem, one for estimating FEM from RGC responses across the retina which negates the need for an extra-retinal mechanism to properly decode spatial information, and another one for making an *optimal* inference about the spatial layout of the stimuli. The presence of a global motion compensation mechanism for FEM was demonstrated by a striking visual illusion (Murakami & Cavanagh, 1998). Surprisingly, when receptive field size and density across the retina and the statistics of FEM under normal viewing conditions are factored in, this model predicted that normal human FEM are not optimal for high acuity tasks (Burak et al., 2010; Pitkow et al., 2007). This theory also predicts that larger stimulus sizes and peripheral cues would improve discrimination at the fovea since estimating FEM would be easier and more accurate in these conditions. It is possible that the absence of optimality might have arisen since the scanning raster was always visible in the present work. Although several lines of evidence against this prediction have been presented (Wehrhahn, 2011), they turned out to be lacking technical precision and proper controls to directly test this prediction (Burak, Rokni, Meister, & Sompolinsky, 2011).

### Covarying factors

We have identified several mediator factors that could explain a significant portion of the variability in the data. Note that the presence of these mediators is not due to the equipment used or stimulus parameters, but reflects the inevitable consequence of foveal presentation of the stimuli. None of these mediators have been reported quantitatively or used to account for data in the previous literature about the roles of FEM. Parafoveal (or peripheral) presentation of stabilized stimuli may not activate all of the aforementioned mediators (e.g., eye ISOA), however, non-foveal presentation of stimuli would defeat the purpose of this study since one cannot make strong inferences about foveal viewing with peripherally presented stimuli. Alternatively, an experiment where stimulus moves in an incongruent manner to avoid chasing can be performed, however, it is unclear whether or not small amplitudes of stimulus motion would still lead to pursuit-like eye movements. The way we chose to address what factors underlie the tuning between performance and gain reported here is to perform a mediation analysis (MacKinnon et al., 2007). This analysis showed that even when retinal motion, eye motion, PRL eccentricity, and microsaccade rate were factored in, gain still had a significant direct effect on performance. This finding suggests that (i) there are additional mediators not considered here, or (ii) “post-retinal” factors such as changes in attentional engagement in the task depending on gain value might be at play.

In order to assign extra-retinal factors a role for perception during FEM, one needs to factor out all possible retinal factors such as retinal ISOA, velocity, acceleration, PRL eccentricity, initial retinal position of the stimuli, etc. Obviously, these factors are not independent from each other, limiting the use of mediation analysis described here. Admittedly, the optimal gain might also be affected by these mediators. A way to compensate for their effects for the purpose of estimating optimal gain might be normalizing performance by each mediator and then testing for tuning. However, this exacerbates the problem since (i) whether a covarying factor is a positive mediator (reducing the effect) or a negative one (increasing the effect) is not known a priori, (ii) the relative contribution of each mediator is different but normalization assumes equal contribution, and (iii) each mediator has a different scale of change across conditions, which could result in numerical instabilities and prevent accurate determination of the optimal gain. Point (iii) can be addressed by log-transforming some mediators (e.g., retinal ISOA) and/or standardizing them, and point (i) can be addressed by using the outcome of a mediation analysis to guide the normalization process, but point (ii) cannot be readily addressed. On the other hand, since visual performance comes about via mediators, there may not be a need for normalizing performance before computing optimal gains. From this perspective, they are not just artifacts to be removed, but the actual underlying factors of visual function. The logic is that whatever the exact value of optimal gain is, visual performance results from an interplay between various mediators, and it may not be possible to uniformly sample the multidimensional space defined by multiple mediators. A case in point, it seems that foveal presentation of a stimulus almost always leads to smooth pursuit-like oculomotor behaviors when the retinal projection of it is stabilized (Poletti et al., 2010).

### Microsaccades

Human retina is non-homogeneous, even within the fovea. Therefore, making microsaccades to redirect gaze to enjoy the highest acuity part of the retina is a reasonable strategy (Cornsweet, 1956; Poletti et al., 2013). Microsaccades are not always initiated voluntarily, however, and recent studies claimed that they often occur after a period of low retinal slip and are executed to avoid fading (Engbert & Mergenthaler, 2006). Their occurrence seems to be coupled to heartbeat as well (Ohl, Wohltat, Kliegl, Pollatos, & Engbert, 2016). Based on the finding that microsaccades cause widespread activity across the visual system and help temporally synchronize neural activity, some researchers supported the view that microsaccades, among other FEM, contribute most to visual function (Martinez-Conde et al., 2013; Masquelier, Portelli, & Kornprobst, 2016; McCamy et al., 2012). In addition, a review of old and new literature on microsaccades led some researchers to conclude that microsaccades do not serve a useful purpose (Collewijn & Kowler, 2008). Nevertheless, in order to address these hypotheses, we performed a series of analyses on microsaccades made by all observers (**Fig. 7**). Since the rate of microsaccades was rather low in our experiments, we combined the data across observers for the following analyses. The low rate of microsaccades itself, especially when retinal image motion was minimized, is an evidence against a primary role for microsaccades for visual processing. In response to partial or complete retinal stabilization for instance, subjects made larger drifts rather than larger or more frequent microsaccades. Moreover, we found evidence for both low retinal slip and gaze redirection, although the evidence for the latter was stronger. More specifically, we found that retinal image velocity was slightly reduced immediately before microsaccade onset, and most microsaccades were made to bring the retinal projection of the stimuli closer to the PRL.

**Figure 7.**
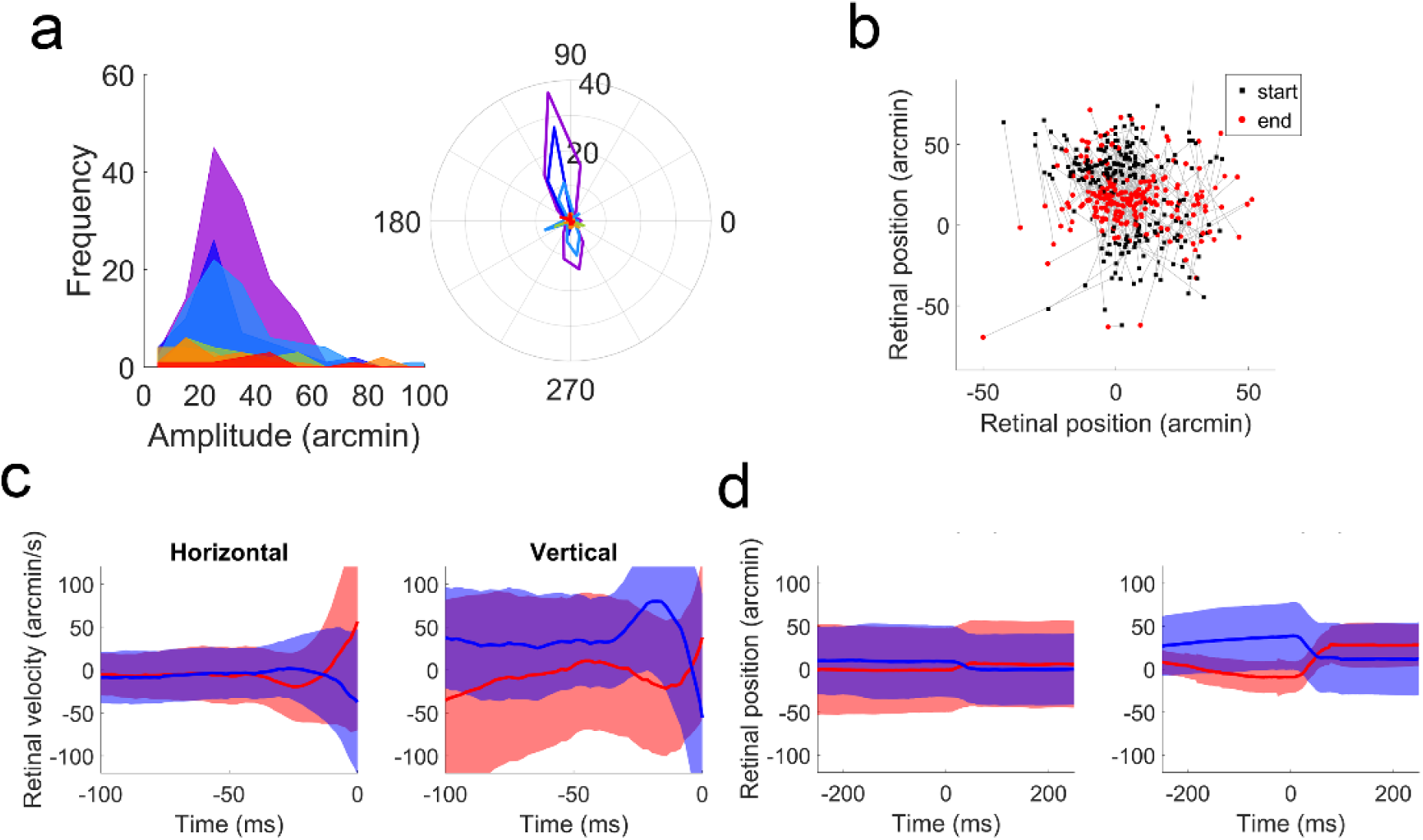
Analyses of microsaccades. **(a)** Amplitude and direction distribution of microsaccades, combined across seven subjects, in Experiment 1. **(b)** The retinal position of the stimuli at the start (black squares) and end (red circles) of microsaccades. Clearly, the primary role of microsaccades was redirecting gaze to compensate for non-homogeneous vision. **(c)** Retinal image velocity just before microsaccades. The blue and red lines represent downward and upward microsaccades (within ±45° from vertical was considered as upward). **(d)** Retinal position of the stimuli across microsaccades. In **(c)** and **(d)**, the panels on the left and right represent data from horizontal and vertical component of the eye movements, respectively.

### Ocular drifts

There are several other facts to be considered when functional roles of drifts and microsaccades are to be determined. First, RGCs are most responsive to light transients, and the time constant of their responses can vary from 30 to 100 ms (Brien, Isayama, Richardson, Berson, & O’Brien, 2002). Second, although the initial burst activity of RGCs in response to a light transient is highly precise, prolonged presentation breaks this temporal synchrony, and the tonic neural activity demonstrates quite a bit of variability (Berry, Warland, & Meister, 1997; Reich, Victor, Knight, Ozaki, & Kaplan, 1997). Encoding spatial information using a rate code with a few spikes necessitates the accumulation of information over time to improve the signal to noise ratio. The presence of FEM makes encoding of spatial information via rate coding even less reliable by further increasing variability in spiking activity. Third, FEM create retinal motion signals that are well beyond motion detection thresholds but not perceived.

From an evolutionary standpoint, it is unclear which of the facts listed so far was the root cause for the others. For instance, whether RGCs prefer light transients and do not respond as strongly after prolonged presentation due to FEM, or FEM exist due to the temporal characteristics of RGC responses is a hard problem to address. In addition, a recent modeling work demonstrated potential alternatives to spatial encoding via rate coding, where FEM do not pose problems to be solved by the visual system, but instead they are part of the solution to efficient information encoding (Ahissar & Arieli, 2012). This also renders the mystery of how the visual system differentiates motion due to FEM from those of external objects a non-issue since if we actually see via FEM, why correct for them? In fact, drifts transform the spectral content of retinal stimulation into spatiotemporal frequencies to which the early visual system is most sensitive (Kuang et al., 2012), but it is unclear whether this is an epiphenomenon or a result targeted by an active and/or adaptive process. However, the current implementation of this model relies on weak assumptions, one of which is that drifts are cyclic (sinusoidal) motions (to drive phase-locking mechanism) within time courses that reflect average fixation duration (∼300 ms). Except in very few instances, we did not observe such patterns (**Fig. 8**).

**Figure 8.**
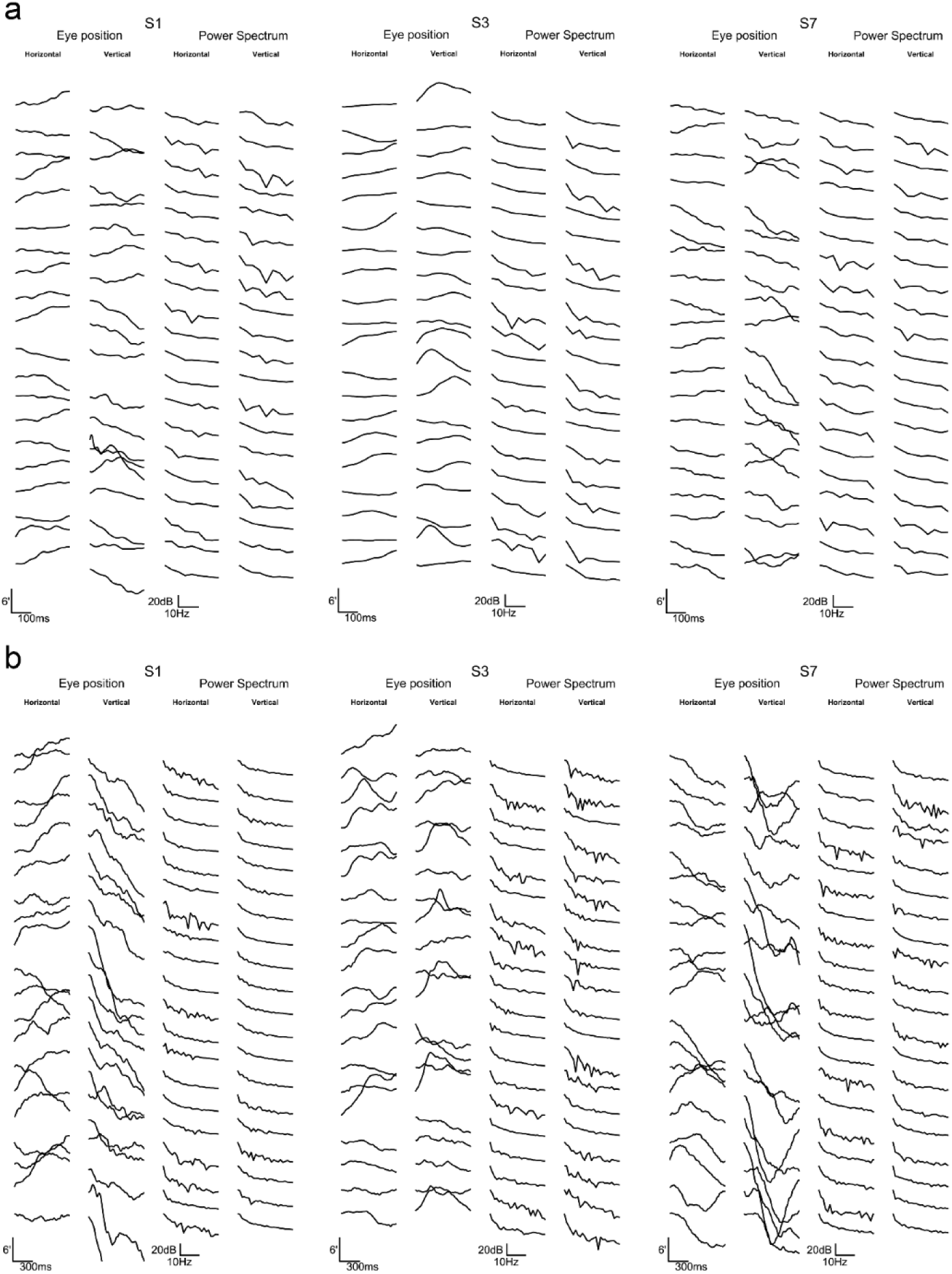
Ocular drifts under normal viewing conditions. To qualitatively test the assumption that drifts are cyclic motions within the time scale of typical fixation (∼300 ms), we randomly sampled 20 eye position traces from three subjects in the zero-gain condition, and computed the power spectra of both the horizontal and vertical components in **(a)** a 300 ms and **(b)** ∼900 ms time windows. If indeed, drifts show three to five cycles per ∼300 ms, this should be visible as clear peaks in the power spectra, and disappear when power spectra are computed over a longer time scale. However, except for a few cases, we did not encounter such motions. For some subjects, much lower-frequency fluctuations were visible, but these fluctuations are too slow to be of any use for fast and efficient temporal encoding. For this particular figure, eye position traces were further filtered with a low-pass filter with a cut-off frequency of 25 Hz. Power spectra are shown with a linear frequency axis with limits from 3 to 30 Hz.

### Abnormal FEM

Some visual/cortical impairments (e.g., amblyopia, central vision loss) result in “abnormal” FEM (Chung, Kumar, Li, & Levi, 2015; Kumar & Chung, 2014). In a computer vision system with limited spatial resolution or blurry optics, it is theoretically possible to achieve “super-resolution” or de-blurring by moving a sensor array. Therefore, we think that to classify FEM as abnormal, one needs to consider several factors such as the amount of blur, receptive field sizes, and contrast sensitivity at the PRL. Otherwise, a genuine strategy of a perfectly normal oculomotor system might be misinterpreted as an artifact. In the case of central vision loss, the use of peripheral PRL leads to changes in all these factors, and it is quite possible that apparently abnormal FEM in these patients might be a way to compensate for these changes. In fact, recent studies on the effects of retinal image motion in peripheral vision reported improvements in reading and discrimination performance with increased motion (Patrick, Roach, & McGraw, 2017; Watson et al., 2012).

### Limitations and future directions

The statistics of FEM may change when a subject's head is restrained compared to head-free viewing (Poletti, Aytekin, & Rucci, 2015). The amplitudes of FEM increase under free viewing. Measurements using a Dual-Purkinje tracker showed that drifts from the two eyes show minimal correlation under head-fixed conditions, and they become mostly conjugate under head-free conditions. However, retinal imaging via a binocular TSLO revealed almost complete conjugacy under head-fixed conditions (Stevenson, Sheehy, & Roorda, 2016). Nonetheless, the conditions reported here may demonstrate a special case of oculomotor control, which need not be optimized since outside the laboratory, we always view the environment with freely moving body and head. In addition, since the stimulus presentation was monocular in the present study, it remains to be seen whether similar tuning functions would be obtained with binocular presentation. It may be that binocular viewing increases the tolerance of the visual system to retinal image motion due to FEM even for high spatial frequencies due to redundancy from the second eye. In addition, as mentioned before, varying retinal image motion while keeping the eye motion unaffected remains to be a challenge. Finally, the different patterns of results in the two experiments reported here also suggest that the relationship between FEM and spatial frequency might be a continuum, from no tuning to optimal tuning with increasing spatiotemporal frequency. Future endeavors along these lines will require denser sampling of the frequency space as well as accurate eye tracking combined with fast stimulus delivery to both eyes.

## Author contributions

MNA and STLC conceived the idea and designed the experiments. MNA and STLC performed the experiments. MNA analyzed the data and wrote the manuscript. CKS, PT, and AR provided technical support for the TSLO system (hardware, electronics, and software), and CKS, PT, AR, and STLC reviewed the manuscript.

## Competing financial interests

MNA and STLC have no financial interest. AR, CKS, and PT hold a patent for the design of the TSLO, and have financial interest in C.Light Technologies.

## Acknowledgements

We would like to thank Jonathan Patrick, Arun Kumar Krishnan and Haluk Ogmen for his comments on the manuscript, and Harold Bedell for many discussions of our results. This study was supported by grants R01-EY012810 (STLC), R01-EY017707 (STLC), R01-EY023591 (AR), P30-EY003176 (core grant) from the National Institutes of Health, and UCSF-CTSI grant TL1-TR001871 (CKS).

